# Plasmodesmal connectivity in C_4_ *Gynandropsis gynandra* is induced by light and dependent on photosynthesis

**DOI:** 10.1101/2022.12.07.519530

**Authors:** Tina B. Schreier, Karin H. Müller, Simona Eicke, Christine Faulkner, Samuel C. Zeeman, Julian M. Hibberd

## Abstract

- In leaves of C_4_ plants the reactions of photosynthesis become restricted between two compartments. Typically, this allows accumulation of C_4_ acids in mesophyll cells and subsequent decarboxylation in the bundle sheath. In C_4_ grasses proliferation of plasmodesmata between these cell types is thought to increase cell-to-cell connectivity to allow efficient metabolite movement. However, it is not known if C_4_ dicotyledons also show this enhanced plasmodesmal connectivity and so whether this is a general requirement for C_4_ photosynthesis is not clear. How mesophyll and bundle sheath cells in C_4_ leaves become highly connected is also not known.
- We investigated these questions using 3D- and 2D- electron microscopy on the C_4_ dicotyledon *Gynandropsis gynandra* as well as phylogenetically close C_3_ relatives.
- The mesophyll-bundle sheath interface of C_4_ *G. gynandra* showed higher plasmodesmal frequency compared with closely related C_3_ species. Formation of these plasmodesmata was induced by light. Pharmacological agents that perturbed chloroplast development or photosynthesis reduced the number of plasmodesmata, but this inhibitory effect could be reversed by the provision of exogenous sucrose.
- We conclude that enhanced formation of plasmodesmata between mesophyll and bundle sheath cells is wired to the induction of photosynthesis in C_4_ *G. gynandra*.

## INTRODUCTION

C_4_ photosynthesis represents a carbon concentrating mechanism that has repeatedly evolved from the ancestral C_3_-type of photosynthesis (Sage et al. (2011)). In leaves of C_4_ plants, HCO_3_^-^ is initially fixed by Phopsho*enol*pyruvate Carboxylase (PEPC) in mesophyll (M) cells into a 4-carbon acid (malate/aspartate). These C_4_ acids then move to bundle sheath (BS) cells for decarboxylation to produce pyruvate and CO_2_. Pyruvate is transferred back to the mesophyll cells where it is reduced to phospho*enol*pyruvate that can accept another HCO_3_^-^ molecule. This spatial separation of carboxylation and decarboxylation between mesophyll and bundle sheath cells builds a high concentration of CO_2_ in bundle sheath cells and in so doing limits the oxygenation side-reaction of RuBisCO (Hatch, 1987). This greatly increases photosynthesis efficiency, particularly in hot and dry environments.

Efficient exchange of metabolites between mesophyll and bundle sheath cells is therefore crucial to the C_4_ pathway and as a consequence compared with the ancestral C_3_ condition C_4_ leaves are typically reconfigured in both biochemistry and structure. Most C_4_ plants have Kranz anatomy - with closely spaced veins and a wreath-like, concentric arrangement of enlarged bundle sheath cells surrounding mesophyll cells that maximises mesophyll-bundle sheath contact sites (Sedelnikova et al., 2018). Kranz anatomy is associated with increased cell-to-cell connectivity between the mesophyll and bundle sheath cells to allow the efficient exchange of metabolites. Metabolite exchange between the two cell types is proposed to occur via passive diffusion through plasmodesmata down a steep concentration gradient of C_4_ metabolites (Hatch, 1987).

Plasmodesmata are regulated channels between adjacent plant cells and diverse in structure: from simple (with single openings in adjacent cells) to complex (highly branched with central cavities), or even asymmetric in their organisation (Ross-Elliott et al., 2017; Faulkner, 2018). Plasmodesmata contain several structural components including a narrow tube of endoplasmic reticulum called the desmotubule, the cytoplasmic sleeve and the plasma membrane (Faulkner, 2018). Plasmodesmata are considered essential for cell-to-cell transport of metabolites in many C_4_ grasses because suberized bundle sheath cell walls likely reduce CO_2_ leakage by blocking apoplastic metabolite transfer (Hatch and Osmond, 1976). Furthermore, C_4_ grasses possess increased numbers of plasmodesmata between mesophyll and bundle sheath cells (Evert et al., 1977; Botha et al., 1992, 1993; Danila et al., 2016). As plasmodesmata occur in clusters (pitfields), increased cell-to-cell connectivity in C_4_ leaves can be a result of increased pit field area or increased numbers of plasmodesmata per pit field area. Danila et al. (2016) observed up to 9-fold increase in plasmodesmal frequency at the mesophyll-bundle sheath interface in C_4_ maize and *Setaria viridis* compared with the C_3_ species rice and wheat. This increase in the C_4_ grasses was due to a 2-fold increase in plasmodesmata numbers per pitfield, and a 5-fold increase in pitfield area. In other C_4_ grasses substantial variation in absolute plasmodesmata frequency was evident but they all possessed greater plasmodesmata frequency than C_3_ species (Danila et al., 2018).

To our knowledge, the distribution of plasmodesmata at the mesophyll-bundle sheath cell interface of C_3_ and C_4_ species has not been studied outside the grasses. Further, the cues that underpin increased plasmodesmata formation are not known. Given the known variation in how increased cell-to-cell connectivity is achieved in C_4_ grasses and the fact that they evolved C_4_ photosynthesis independently from C_4_ dicotyledenous lineages, we assessed plasmodesmata distribution in leaves of C_3_ *Tarenaya hassleriana* and C_4_ *Gynandropsis gynandra* that both belong to the *Cleomaceae* (Brown et al., 2005; Marshall et al., 2007) which is sister to the *Brassicaceae. G. gynandra* has been developed as a C_4_ model (Brown et al., 2005; Marshall et al., 2007; Koteyeva et al., 2011; Bräutigam et al., 2011). We discovered that plasmodesmal frequency is up to 8-fold higher at the mesophyll-bundle sheath cell interface in mature leaves of C_4_ *G. gynandra* compared with that in C_3_ species. Moreover, these increased numbers of plasmodesmata are rapidly established during deetiolation. Pharmacological studies using multiple chloroplast inhibitors demonstrated that light, functional chloroplasts and photosynthesis are required to initiate plasmodesmata formation at mesophyll-bundle sheath cell interface of *G. gynandra*. Provision of exogenous sucrose can rescue defects in chloroplasts and photosynthesis. We conclude that increased plasmodesmatal connection is likely an unifying feature of all two-celled C_4_ plants, and that during the evolution of the C_4_ pathway the increased formation of secondary plasmodesmata is induced by the induction of photosynthesis itself.

## MATERIAL AND METHODS

### Plant Material and growth conditions

*G. gynandra* and *T. hassleriana* seeds were germinated on wet filter papers in petri dishes. For *G. gynandra*, germination was initiated by exposing seeds to 30°C for 24 h. For *T. hassleriana*, germination was stimulated by an alternating temperature regime of 12 h 32°C then 12 h at 20°C for 5 consecutive days. After germination, *G. gynandra* and *T. hassleriana* seedlings were planted in 10:1 ratio of M3 compost (Levington Advance, Pot and Bedding, High Nutrient):fine vermiculite in individual pots. *A. thaliana* (Col-0) was sown onto potting compost (Levington Advance, Solutions) with 0.17 g L^-1^ insecticide (thiacloprid, Exemptor) and stratified for 48 h at 4°C. Around 2 weeks after germination, individual seedlings were transplanted to individual pots.

To sample of mature leaves, plants were grown in a climate-controlled growth chamber with 16-h light and 8-h dark. *G. gynandra* and *T. hassleriana* were grown at 350 μmol photons m^-2^ s^-1^ at 25°C with a 60% (v/v) relative humidity and ambient CO_2_. *A. thaliana* plants were grown under identical conditions except light intensity was 150 μmol photons m^-2^ s^-1^. All plants were watered by an automated system whereby the bottom of the trays was flooded to a depth of 4 cm every 48 h for 10 min, after which the irrigation water was drained.

For deetiolation experiments, *G. gynandra* seeds were germinated with the addition of 0.15% (v/v) plant preservative mixture (Apollo Scientific, CAS: 26172-55-4) to the wet filter paper. Germinated seedlings were transferred to square plates containing half-strength MS (Murashige and Skoog) salts with B5 vitamins (Duchefa Biochemie BV) and 0.8% (w/v) agar (Melford) in the dark. Plates were grown in the plant growth cabinet (Panasonic MLR-352 PE) at 20 °C with continuous light intensity of 150 μmol m^-2^ s^-1^. Plates were covered with aluminum foil for three consecutive days to ensure no light was able to penetrate. Aluminum foil was removed on day 3 and to allow de-etiolation plants grown for an additional 24 to 48 h in the light. For sucrose supplementation, 10 g L^-1^ sucrose was added to the half-strength MS media. For inhibitor treatments, 500 μM lincomycin (Sigma Aldrich), 50 μM norflurazon (Sigma Aldrich) and 20 μM DCMU (Sigma Aldrich) were added to the half-strength MS media before the media was poured in the individual petri dishes. As norflurazon and lincomycin were dissolved in ethanol, the control and DCMU treatments included an equivalent amount of ethanol in the media.

### Electron microscopy

Samples from 5-8 individual seedlings at each time point were harvested for electron microscopy. Leaf segments (~2 mm^2^) were excised with a razor blade and immediately fixed in 2% (v/v) glutaraldehyde and 2% (w/v) formaldehyde in 0.05 - 0.1 M sodium cacodylate (NaCac) buffer (pH 7.4) containing 2 mM calcium chloride. Samples were vacuum infiltrated overnight, washed 5 times in 0.05 – 0.1 M NaCac buffer, and post-fixed in 1% (v/v) aqueous osmium tetroxide, 1.5% (w/v) potassium ferricyanide in 0.05 M NaCac buffer for 3 days at 4°C. After osmication, samples were washed 5 times in deionized water and post-fixed in 0.1% (w/v) thiocarbohydrazide for 20 min at room temperature in the dark. Samples were then washed 5 times in deionized water and osmicated for a second time for 1 h in 2% (v/v) aqueous osmium tetroxide at room temperature. Samples were washed 5 times in deionized water and subsequently stained in 2% (w/v) uranyl acetate in 0.05 M maleate buffer (pH 5.5) for 3 days at 4°C, and washed 5 times afterwards in deionized water. Samples were then dehydrated in an ethanol series, transferred to acetone, and then to acetonitrile. Leaf samples were embedded in Quetol 651 resin mix (TAAB Laboratories Equipment Ltd) and cured at 60°C for 2 days.

### Transmission electron microscopy (TEM) and Scanning electron microscopy (SEM)

For TEM, ultra-thin sections were cut with a diamond knife using a Leica Ultracut microtome and collected on copper grids and examined in a FEI Tecnai G2 transmission electron microscope (200 keV, 20 μm objective aperture). Images were obtained with an AMT CCD camera. For SEM of plasmodesmata pitfields in *G. gynandra, T. hassleriana* and *A. thaliana*, samples were prepared according to Danila et al. (2018). In summary, mature leaves were cut into 10-20 mm strips and fixed in 2% (v/v) glutaraldehyde and 2% (w/v) formaldehyde in 0.05 - 0.1 M sodium cacodylate (NaCac) buffer (pH 7.4) containing 2 mM calcium chloride under vacuum infiltration overnight at RT. Leaf tissue was dehydrated in an ethanol series and critical point dried (CPD) in a Quorum E3100 dryer. CPD leaf samples were ripped apart using forceps and sticky tape. Ripped samples were mounted on aluminum SEM stubs using conductive carbon tabs (TAAB), sputter-coated with a thin layer of iridium (15 nm) and imaged in a Verios 460 scanning electron microscope (FEI, Hillsboro, OR) run at an accelerating voltage of 2 keV and 25 pA probe current. Low magnification images were aquired with an Everhart-Thornley detector whilst high-resolution images were acquired using the through-lens detector in immersion mode.

For 2D SEM mapping, ultra-thin sections were placed on Melinex (TAAB Laboratories Equipment Ltd) plastic coverslips mounted on aluminum SEM stubs using conductive carbon tabs (TAAB Laboratories Equipment Ltd), sputter-coated with a thin layer of carbon (~ 30 nm) to avoid charging and imaged in a Verios 460 scanning electron microscope at 4 keV accelerating voltage and 0.2 nA probe current using the concentric backscatter detector in field-free (low magnification) or immersion (high magnification) mode (working distance 3.5 – 4 mm, dwell time 3 μs, 1536 x 1024 pixel resolution). For plasmodesmata frequency quantification, SEM stitched maps were acquired at 10,000X magnification using the FEI MAPS automated acquisition software. Greyscale contrast of the images were inverted to allow easier visualisation.

Serial block face scanning electron microscopy (SBF-SEM) was performed on Quetol 651 resin-embedded mature leaf samples of *G. gynandra, T. hassleriana* and *A. thaliana* as described above. Overviews of leaf cross-sections and the zoomed stacks of the mesophyll – bundle sheath cell interface (≈300-400 images) were acquired through sequentially sectioning the block faces at 50 nm increments and imaging the resulting block-face by SEM. Images were acquired with a scanning electron microscope TFS Quanta 250 3VIEW (FEI, Hillsboro, OR) at 1.8-2 keV with an integrated 3VIEW stage and a backscattered electron detector (Gatan Inc., Pleasanton, CA, USA). Images were aligned and smoothed using the plugins MultiStackReg and 3D median filter on ImageJ.

Plasmodesmal frequency from 2D and 3D EM images was determined using published methods (Koteyeva et al., 2014; Botha, 1992). Briefly, plasmodesmal frequency was determined as the number of plasmodesmata observed per μm of length of shared cell interface between two cell types (mesophyll – bundle sheath, mesophyll – mesophyll, bundle sheath – bundle sheath). Plasmodesmata numbers and cell lengths were determined using ImageJ software. Plasmodesmata were defined as dark channels in the EM images. Depending on plasmodesmata orientation, the entire channel was sometimes not visible on 2D EM images, and so only channels that spanned more than half of the cell wall width were counted.

### Chlorophyll fluorescence measurement

Chlorophyll fluorescence measurements were carried out using a CF imager (Technologica Ltd, UK) and image processing software provided by the manufacturer. Seedlings were placed in the dark for 20 min evaluate dark-adapted minimum fluorescence (*Fo*), dark-adapted maximum fluorescence (*Fm*) and then variable fluorescence *Fv* (*Fv* = *Fm*–*Fo*). All chlorophyll fluorescence images of inhibitor-treated seedlings within each experiment were acquired at the same time in a single image, measuring a total of 8 seedlings per treatment.

### Statistical analysis

In violin plots, the middle line represents the median, the box and whiskers represent the 25 to 75 percentile and minimum-maximum distributions of the data. Letters show the statistical ranking using a one-way ANOVA and post hoc Tukey test (different letters indicate differences at P<0.05). Values indicated by the same letter are not statistically different. Data was analyzed using RStudio 2022.07.2+576.

## RESULTS

### Plasmodesmata frequency is higher in C_4_ *G. gynandra* leaves compared with C_3_ *A. thaliana* and *T. hassleriana*

We first explored whether the increased plasmodesmal connectivity between mesophyll and bundle sheath cells found in C_4_ grasses was also present in the C_4_ dicotyledon *Gynandropsis gynandra*. Transmission electron microscopy was used to examine the mesophyll-bundle sheath cell interface in mature leaves of *G. gynandra* and the closely related C_3_ species *Tarenaya hassleriana* (also a member of the *Cleomaceae*) as well as C_3_ *Arabidopsis thaliana*. Plasmodesmata were more abundant between mesophyll and bundle cells in C_4_ *G. gynandra* compared with both C_3_ species (Fig. **1**). Increased physical connectivity was specific to this interface, and no obvious increases were detected at the mesophyll-mesophyll or bundle sheath-bundle sheath cell interfaces in any species (Supporting Information Fig. **S1**).

**Figure 1.**
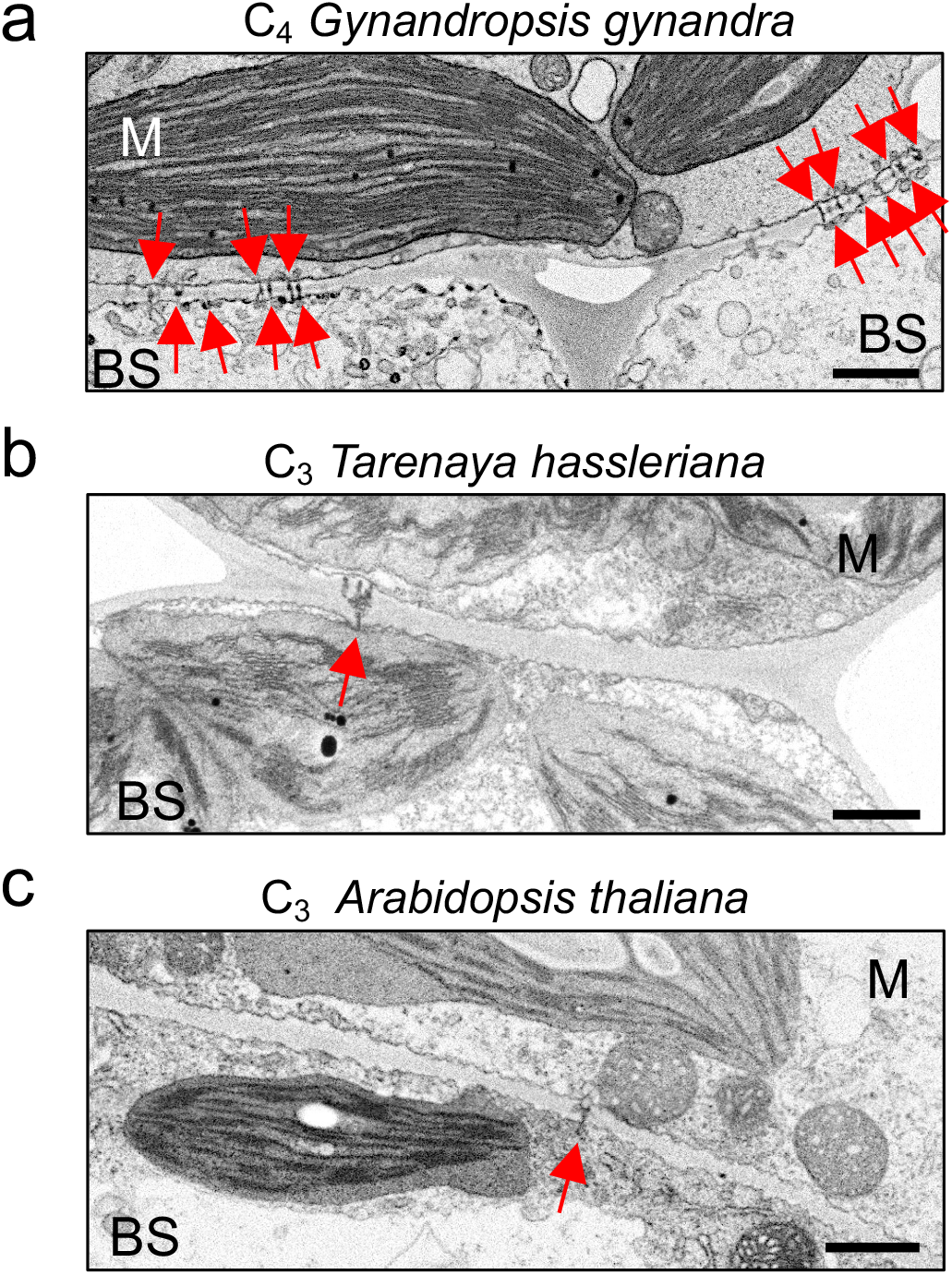
The Mesophyll (M) - Bundle Sheath (BS) cell interface of C_4_ *Gynandropsis gynandra* has more plasmodesmata than closely related C_3_ species. Representative transmission electron micrographs of M-BS interfaces in (**a**) C_4_ *G. gynandra*, (**b**) C_3_ *Tarenaya hassleriana* and (**c**) C_3_ *Arabidopsis thaliana*. Mature leaves were harvested from 4-week-old *G. gynandra* and *T*. hassleriana, or 3-week-old *A. thaliana* plants. Red arrows indicate individual plasmodesma. Scale bar represents 1 μm.

To quantify plasmodesmata numbers between mesophyll and bundle sheath cells, we conducted serial block-face scanning electron microscopy (SBF-SEM). SBF-SEM offers excellent resolution in 3D and has previously been used to quantify plasmodesmata in other systems (Ross-Elliott et al., 2017; Paterlini and Belevich, 2022). Thin sections prepared from fully expanded true leaves of *G. gynandra, T. hassleriana* and *A. thaliana* were imaged, and an area of the mesophyll-bundle sheath cell interface identified for serial block face sectioning (Fig. **2a-c**). From each species, between 281-438 serial transverse sections per mesophyll-bundle sheath cell interface were collected and compiled into videos (Supporting Information Videos **S1-3**). Using these SBF-SEM sections we quantified plasmodesmata frequency by determining the number of plasmodesmata per length of mesophyll-bundle sheath cell interface imaged in 3D (Fig. **2d**). In C_4_ *G. gynandra*, plasmodesmata were visible in almost every mesophyll-bundle sheath cell interface assessed such that only 20 out of 467 contained no plasmodesmata (Fig. **2d**). In contrast, in the two C_3_ species plasmodesmata were not detected in the majority of interfaces (263/367 for *T. hassleriana*, 628/886 for *A. thaliana*). Because plasmodesmata appear in clusters (pitfields) rather than being equally distributed, a wide range of plasmodesmal frequencies per section were observed between mesophyll and bundle sheath cells in all three species. However, there were more sections with higher frequencies observed at the mesophyll-bundle sheath interface of C_4_ *G. gynandra*, and this resulted in a 13-fold increase in the mean frequency compared with C_3_ *T. hassleriana* and C_3_ *A. thaliana* (Fig. **2d**). Plasmodesmal frequencies between mesophyll and bundle sheath cells of the C_3_ species *T. hassleriana* and *A. thaliana* were not significantly different to each other and were low compared with C_4_ *G. gynandra*.

**Figure 2.**
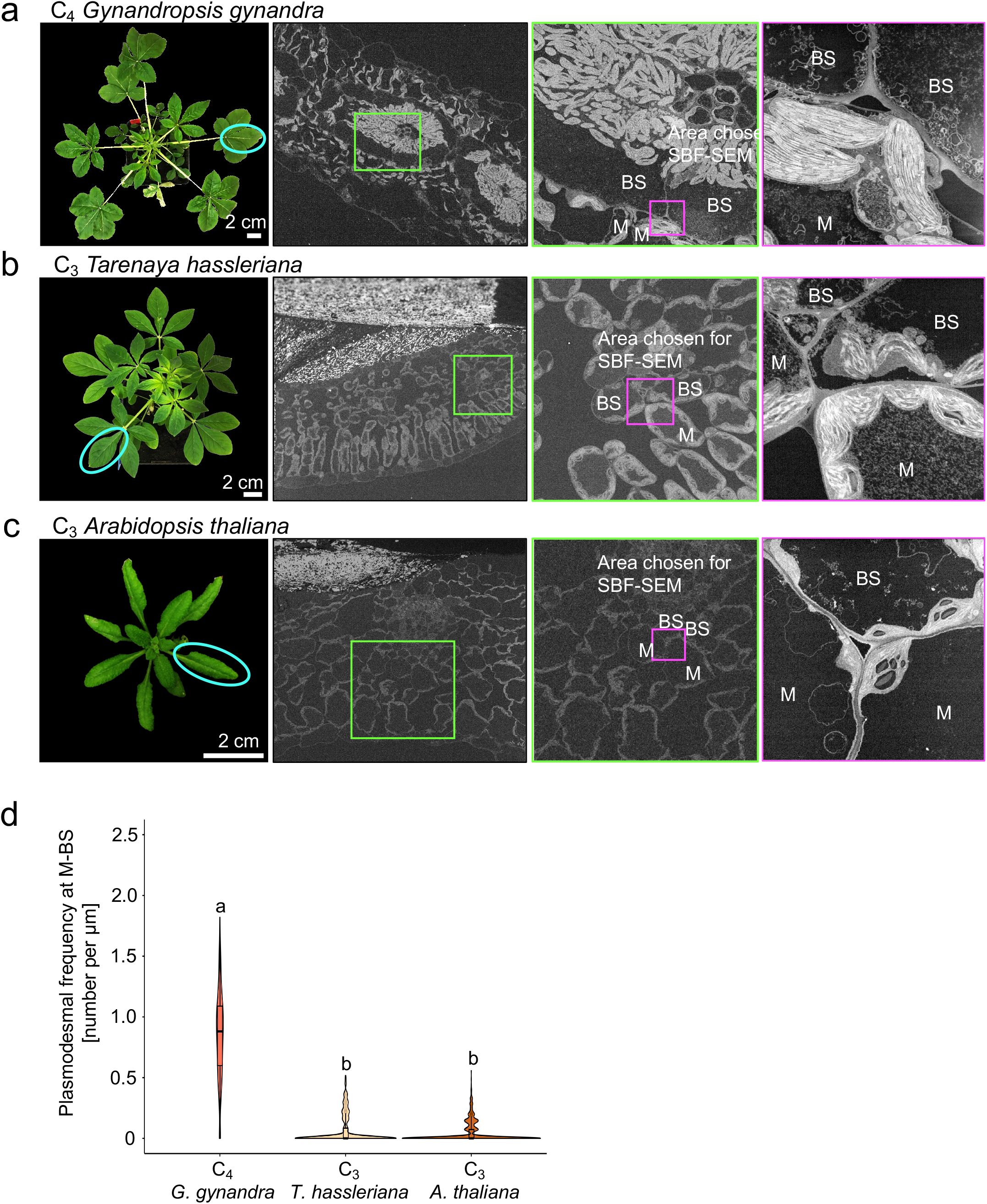
3D Serial Block Face-SEM (SBF-SEM) analysis of plasmodesmata number at the Mesophyll (M) - Bundle Sheath (BS) cell interface. Left panels: Photographs of 4-week-old (**a**) C_4_ *G. gynandra* and (**b**) C_3_ *T. hassleriana*, and 3-week- old (**c**) C_3_ *A. thaliana* plants. Mature leaves harvested for plasmodesmata quantification are circled. Mid left panels: Representative scanning electron micrographs of leaf cross sections from **(a)** C_4_ *G. gynandra*, **(b)** C_3_ *T. hassleriana* and (**c**) C_3_ *A. thaliana*. Mid right panels: Zoomed in image of the region marked by a green box showing area of M-BS cell interface used for SBF-SEM analysis (magenta). Left panels: Single frame of compiled SBF-SEM data into Supporting Information Videos **S1-3** of (**a**) C_4_ *G. gynandra* (**b**) C_3_ *T. hassleriana* and (**c**) C_3_ *A. thaliana*. (**d**) Violin plot of plasmodesmal frequencies measured at M-BS cell interfaces in the three plant species using 3D SBF-SEM data. As some sections contained more than one M-BS cell interface, plasmodesmata frequencies were quantified from a total of 476 individual M-BS interfaces for *G. gynandra*, 367 for *T. hassleriana* and 886 for *A. thaliana*. Box and whiskers represent the 25 to 75 percentile and minimum-maximum distributions of the data. Letters show statistical ranking using a *post hoc* Tukey test (with different letters indicating statistically significant differences at P<0.05). Values indicated by the same letter are not statistically different.

SBF-SEM provides an excellent 3D view of plasmodesmata frequency and distribution but is relatively low throughput and so limited numbers of cell interfaces can be visualised per unit time. We therefore used 2D electron microscopy to further explore the high occurrence of plasmodesmata at the mesophyll-bundle sheath cell interface of C_4_ *G. gynandra*. Large areas of leaf sections were imaged at high resolution using 2D Scanning Electron Microscopy (SEM) mapping such that automated serial imaging at 10,000X magnification and subsequent image stitching enabled visualization of plasmodesmata at numerous interfaces of the same 2D section (Fig. **3a**). Representative SEM maps in which cell interfaces (mesophyll-bundle sheath, mesophyll-mesophyll and bundle sheath-bundle sheath) were pseudocoloured according to plasmodesmal frequency, and consistent with the 3D SBF-SEM analysis reported above, illustrated that plasmodesmata were specifically enriched at the mesophyll-bundle sheath interface of C_4_ *G. gynandra* (indicated by the numerous green-coloured interfaces). In contrast, frequency was lower and more uniform between all cellular interfaces in the C_3_ species (indicated by the pink and orange pseudocoloured cell interfaces) (Fig. **3a**). Plasmodesmata frequencies were quantified from at least three SEM maps originating from three independent plants (biological replicates; 10-40 individual mesophyll – bundle sheath, mesophyll-mesophyll and bundle sheath-bundle sheath cell interfaces per biological replicate). This showed that plasmodesmata numbers between mesophyll and bundle sheath cells were more than 8-fold higher in C_4_ *G. gynandra* compared with both C_3_ species. The three cellular interfaces (mesophyll-bundle sheath, mesophyll-mesophyll and bundle sheath-bundle sheath) in both C_3_ species had similar plasmodesmal frequencies. Interestingly, plasmodesmal frequency of all three types of cell interface in *G. gynandra* was significantly higher than that of the corresponding interface in each of the two C_3_ species. For example, the mesophyll-mesophyll and bundle sheath-bundle sheath interfaces were approximately 3 to 4-fold and 2-fold higher in C_4_ *G. gynandra* compared with *T. hassleriana* and *A. thaliana* respectively indicating that cell-to-cell connectivity is generally enhanced between photosynthetic cells of the C_4_ species (Fig. **2b**). Plasmodesmata frequencies estimated from analysis of numerous mesophyll-bundle sheath cell interfaces using this 2D SEM mapping were not statistically different from the frequencies obtained from multiple serial sections of the mesophyll-bundle sheath interface using SBF-SEM (Supporting Information Fig. 2). To allow greater replication and sampling subsequent analysis was therefore carried out with the 2D SEM mapping technique.

**Figure 3.**
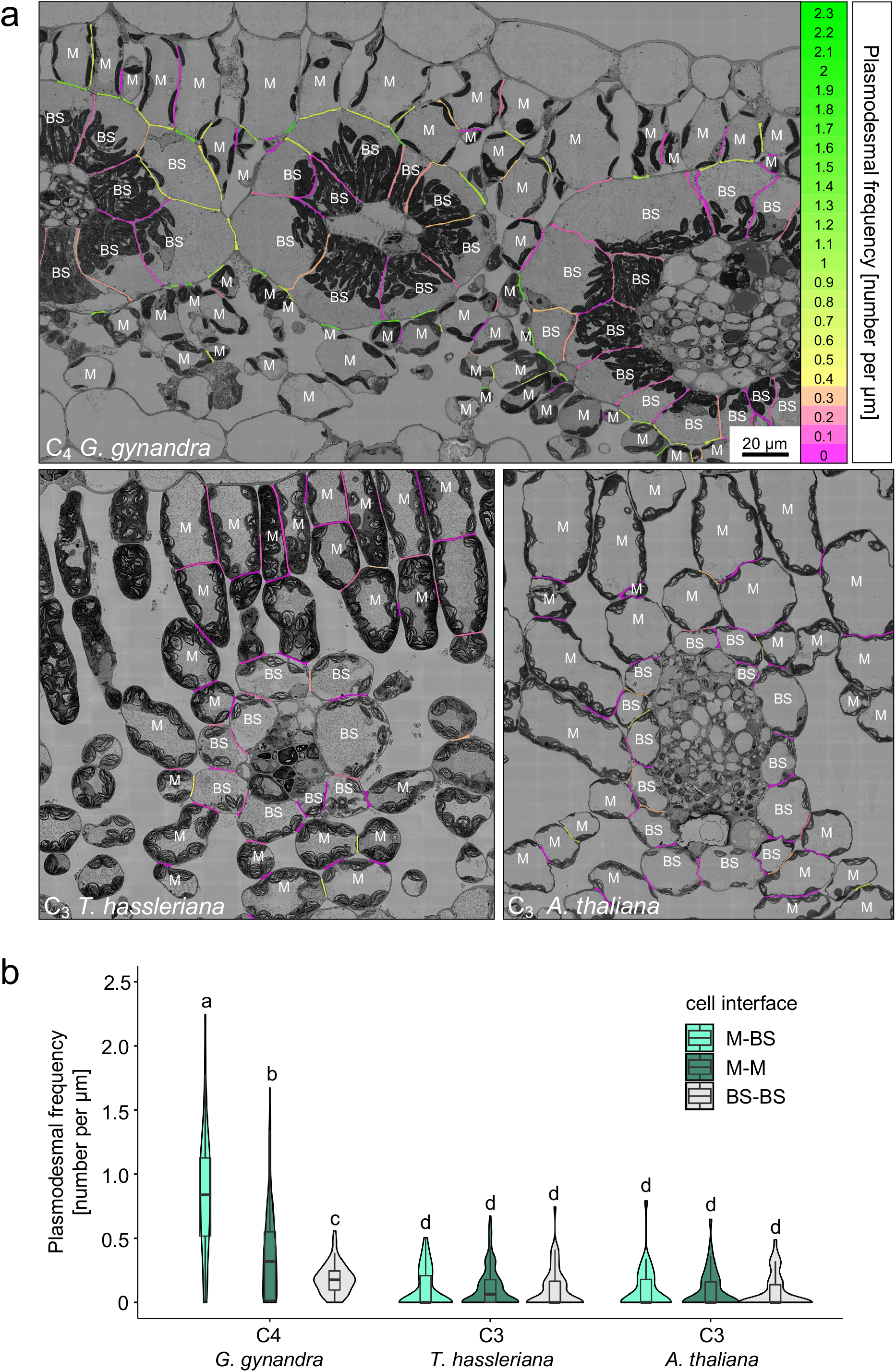
Plasmodesmata frequency in C_4_ *G. gynandra* is higher between M-BS cells compared with other interfaces. **(a)** Heatmap of plasmodesmata distribution. Cell interfaces in high-resolution 2D SEM maps of C_4_ *G. gynandra*, C_3_ *T. hassleriana* and C_3_ *A. thaliana* leaf cross sections were coloured according to plasmodesmal frequency (number of plasmodesmata observed on the interface, divided by the interface length [μm]). (**b**) Plasmodesmal frequency for M-BS, M-M, and BS-BS interfaces in *G. gynandra, T. hassleriana* and *A. thaliana* mature leaves quantified using high-resolution 2D SEM maps. For *G. gynandra, n* = 86 M-M, *n* = 96 M-BS and *n* = 70 BS-BS cell interfaces were quantified. For *T. hassleriana, n* = 202 M-M, *n* = 80 M-BS and *n* = 77 BS-BS cell interfaces were quantified. For *A. thaliana, n* = 45 M-M, *n* = 37 M-BS and *n* = 54 BS-BS cell interfaces were quantified. All interfaces were quantified from leaf samples of at least three individual plants (biological replicates) per species. Box and whiskers represent the 25 to 75 percentile and minimummaximum distributions of the data. Letters show the statistical ranking using a *post hoc* Tukey test (different letters indicate statistically significant differences at P < 0.05). Values indicated by the same letter are not statistically different.

To investigate the relationship between increased frequency of plasmodesmata at the mesophyll-bundle sheath interface and pit fields, we visualized pitfields using SEM by tearing critical point dried mature leaves as described in Danila et al. (2016). Pitfields were clearly visible at the mesophyll-bundle sheath interface in all species, but unlike the previous work in grasses individual plasmodesmata within the pitfields could not be distinguished (Supporting Information Fig. **3a**). When we measured the mean area of pitfields in each species there was no clear difference. This suggests that the increased plasmodesmata frequency at mesophyll-bundle sheath in *G. gynandra* most likely results from increased pit field numbers per cell interface rather than enlarged pit fields that contain more plasmodesmata (Supporting Information Fig. **3b**).

### Increased plasmodesmal frequency between mesophyll and bundle sheath cells of C_4_ *G. gynandra* is established after exposure to light

Induction of the photosynthetic apparatus associated with the C_4_ pathway, such as chloroplast development and C_4_ gene expression typically occurs rapidly in response to light (Shen et al., 2009; Singh et al., 2021). Such de-etiolation analysis is simplest if cotyledons can be analysed. As cotyledons of *G. gynandra* have C_4_ anatomy (Koteyeva et al., 2011) we next examined plasmodesmata in this tissue during de-etiolation. Cross sections of cotyledons showed that Kranz anatomy was already partially developed in 3-day-old dark grown seedlings (Fig. **4a**). For example, veins were closely spaced, and bundle sheath cells contained abundant organelles. However, after 24 h of light cotyledons had almost doubled in size and substantial cell expansion and formation of air spaces was evident (Fig. **4a**). High-resolution 2D SEM maps from cross sections of at least three cotyledons (biological replicates) of *G. gynandra* were obtained at 0 h, 24 h and 48 h after transfer to light. In dark-grown seedlings plasmodesmal frequency at mesophyll-bundle sheath, mesophyll-mesophyll, and bundle sheath-bundle sheath were similar (n = 204) (Fig. **4c,d**). However, after light induction plasmodesmal frequency increased 1.7-fold after 24 h and 2.5-fold after 48 h between mesophyll and bundle sheath cells of *G. gynandra* (Fig. **4b-d**). There was also a small increase in plasmodesmata numbers between mesophyll cells after light exposure. These responses were specific to de-etiolation because growth in the dark for 48 h did not increase plasmodesmata numbers (Supporting Information Fig. **4a-d**). These data indicate that as with true leaves, cotyledons of *G. gynandra* develop high plasmodesmal connectivity between mesophyll and bundle sheath cells, and that this takes place rapidly in response to light. We conclude that light is a crucial developmental cue for the formation of secondary plasmodesmata at the mesophyll-bundle sheath interface in the C_4_ plant *G. gynandra*.

**Figure 4.**
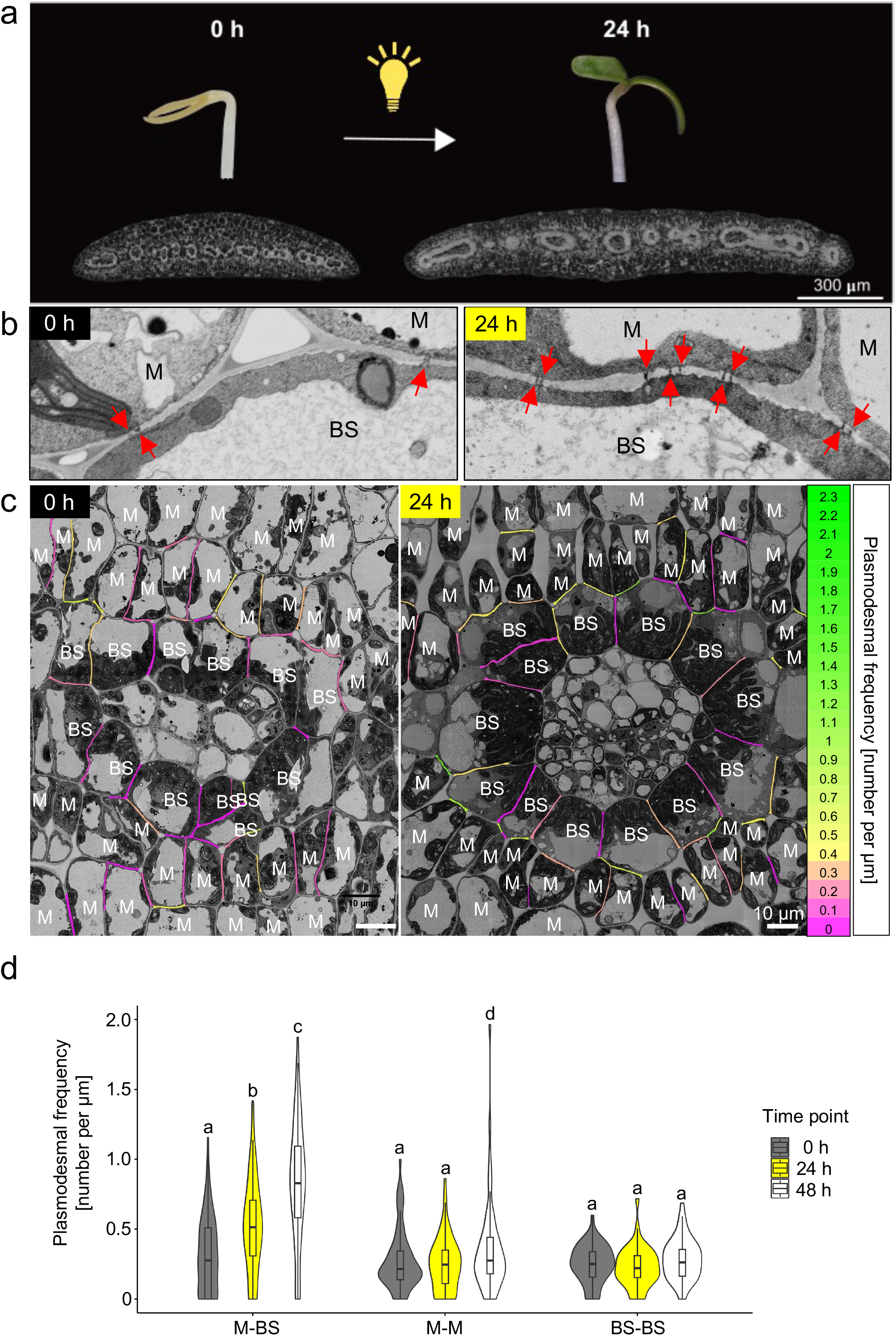
Light acts as a developmental cue to increase plasmodesmata formation at the M-BS cell interface in cotyledons of C_4_ *G. gynandra*. **(a)** Photographs of representative etiolated (left) and deetiolated (right) *G. gynandra* seedlings and scanning electron micrographs of cotyledon cross sections at 0 h and 24 h after light. **(b)** Representative scanning electron micrographs of M-BS interfaces in C_4_ *G. gynandra* cotyledons. Red arrows indicate individual plasmodesma. Scale bar = 1 μm **(c)** Heatmap of plasmodesmata distribution. Cell interfaces in high-resolution 2D-SEM maps of C_4_ *G. gynandra* cotyledon cross sections, harvested prior to light induction (0 h time point) and after light (24 h time point) were coloured according to plasmodesmal frequency (number of plasmodesmata observed on the interface, divided by the interface length [μm]). **(d)** Plasmodesmata frequency per μm cell interfaces (M-BS, M-M, BS-BS) in *G. gynandra* cotyledons was quantified during dark to light transition (0 h, 24 h and 48 h time point) using high resolution 2D SEM maps. For the 0 h time point, *n* = 81 (M-BS), *n* = 74 (M-M) and *n* = 49 (BS-BS) cell interfaces were quantified. For the 24 h time point, *n* = 69 (M-BS), *n* = 70 (M-M) and *n* = 42 (BS-BS) cell interfaces were quantified. For the 48h time point, *n* = 90 (M-BS), *n* =60 (M-M) and *n* = 49 (BS-BS) cell interfaces were quantified. All interfaces were quantified from cotyledon samples of at least 3 individual seedlings (biological replicates) per time point. The box and whiskers represent the 25 to 75 percentile and minimum-maximum distributions of the data. Letters show the statistical ranking using a one-way ANOVA with a *post hoc* Tukey test (different letters indicate statistically significant differences at P < 0.05). Values indicated by the same letter are not statistically different.

### Functional chloroplasts are required for light-induced formation of plasmodesmata between the mesophyll and bundle sheath

De-etiolation involves the transition from skotomorphogenic to photomorphogenic growth whereby fully photosynthetic chloroplasts develop from etioplasts within hours of light exposure (Pipitone et al., 2021; Singh et al., 2021; Cackett et al., 2021). Therefore, it is possible that the increase in plasmodesmal connectivity between mesophyll and bundle sheath cells during de-etiolation is either a direct response to light or is triggered by signals from the chloroplast or photosynthesis. To investigate this, we used inhibitors with distinct modes of action to perturb chloroplast function. Lincomycin and norflurazon block plastid translation and carotenoid biosynthesis respectively and so stop the development of chloroplasts from etioplasts (Mulo et al., 2003; Chamovitz et al., 1991); Fig. **5b**). DCMU [3-(3,4-dichlorophenyl)-1,1-dimethylurea] blocks the electron transport chain at Photosystem II (PSII) (Trebst, 2007) and thus inhibits photosynthesis directly. Seedlings were grown with and without each inhibitor and transferred to light for 48 h. Lincomycin- and DCMU-treated seedlings had pale yellow cotyledons indistinguishable from non-treated controls. Norflurazon treatment generated seedlings with white cotyledons, consistent with compromised carotenoid accumulation (Fig. **5a**). Etioplast ultrastructure was largely unaffected by the inhibitors (Fig. **5b**). After 48 h of light cotyledons of controls and DCMU-treated seedlings were green and etioplasts had developed into chloroplasts (Fig. **5a,b**). Norflurazon and lincomycin-treated seedlings had pale cotyledons even after light induction and the etioplast-to-chloroplast development was arrested (Fig. **5a,b**). To confirm that each inhibitor had the expected effect on chloroplast function we used chlorophyll fluorescence imaging to quantify *F_v_/F_m_* which provides a read-out for the maximum quantum efficiency of Photosystem II. Each inhibitor drastically reduced *F_v_/F_m_* compared with controls (Fig. **5c,d**). Norflurazon-treated seedlings were not visible on the chlorophyll fluorescence imager as chlorophyll content was too low.

**Figure 5.**
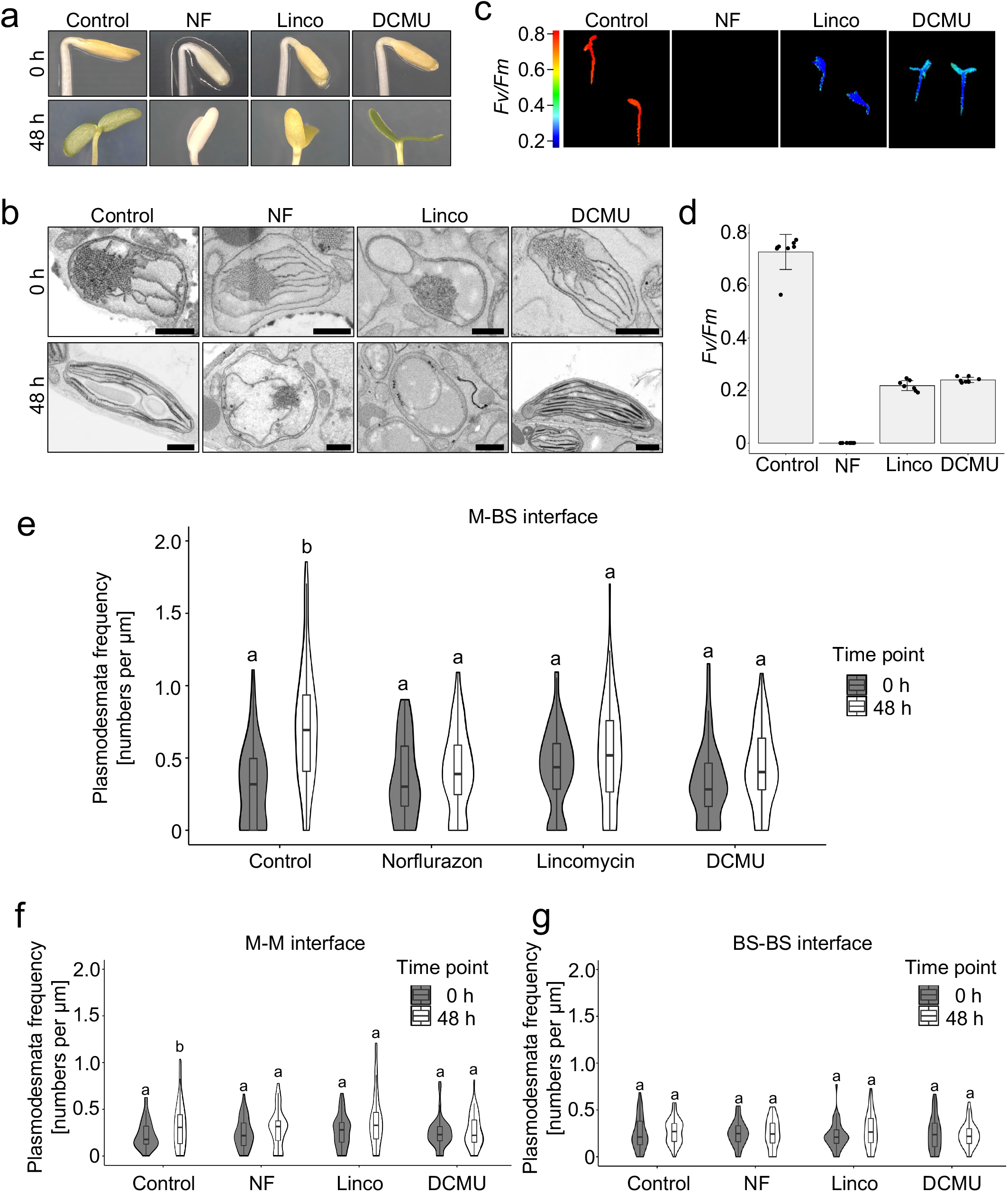
Inhibitors of chloroplast function reduce plasmodesmata formation at the M-BS interface. The effect of norflurazon (NF), lincomycin (Linco) and DCMU were tested. **(a)** Photographs of *G. gynandra* seedlings treated with inhibitors during deetiolation at 0 h and 48 h. (**b**) Scanning electron micrographs of etioplasts (0 h) and mature chloroplasts (48 h) in inhibitor-treated and untreated (Control) *G. gynandra* seedlings. Scale bar represents 1 μm **(c)** Chlorophyll fluorescence images of maximum quantum efficiency of PSII photochemistry (*Fv/Fm*) from 48 h deetiolated *G. gynandra* seedlings treated with NF, Linco and DCMU, as well as untreated seedlings (Control). **(d)** *Fv/Fm* measured in inhibitor-treated and untreated *G. gynandra* seedlings at 48 h after light induction. Bars represent mean ± standard deviation from *n* = 7-8 individual seedlings, dots represent individual data points. (**e-g**) Plasmodesmata frequency per μm cell interfaces in *G. gynandra* cotyledons was quantified during dark to light transition (0 h and 48 h time point) for each individual inhibitor treatment using high-resolution 2D SEM maps: (**e**) M-BS, (**f**) M-M and (**g**) BS-BS. (**e**) For M-BS interface: 0h control *n* = 59, 0h norflurazon *n* = 53, 0h lincomycin *n* = 55, 0h DCMU *n* = 50, 48h control *n* = 85, 48h norflurazon *n* = 66, 48h lincomycin *n* = 90, 48h DCMU *n* = 50 cell interfaces were quantified. (**f**) For M-M interface: 0h control *n* = 41, 0h norflurazon *n* = 43, 0h lincomycin *n* = 45, 0h DCMU *n* = 41, 48h control *n* = 45, 48h norflurazon *n* = 45, 48h lincomycin *n* = 45, 48h DCMU *n* = 45 cell interfaces were quantified. (**g**) For BS-BS interface: 0h control *n* = 41, 0h norflurazon *n* = 39, 0h lincomycin *n* = 44, 0h DCMU *n* = 38, 48h control *n* = 45, 48h norflurazon *n* = 45, 48h lincomycin *n* = 43, 48h DCMU *n* = 45 cell interfaces were quantified. All interfaces were quantified from cotyledon samples of at least 3 individual seedlings (biological replicates) per time point. The box and whiskers represent the 25 to 75 percentile and minimum-maximum distributions of the data. Letters show the statistical ranking, pairwise comparison of 0h and 48 h time point for each treatment, using a *post hoc* Tukey test (different letters indicate statistically significant differences at P < 0.05). Values indicated by the same letter are not statistically different.

Using 2D SEM maps we quantified plasmodesmal frequency in nearly 1200 independent cell interfaces (549 interfaces for the 0 h time point, 649 interfaces for the 48 h time point). None of the three inhibitors affected plasmodesmal frequency at any cell interface in dark-grown seedlings (Fig. **5e-g**). However, despite cotyledon expansion being unaffected by the inhibitors during de-etiolation (Supporting Information Fig. **5a**) plasmodesmal frequencies did not increase significantly in seedlings treated with norflurazon, lincomycin or DCMU (Fig. **5e-g**, Supporting Information Fig. **5b**). In summary, inhibitors that perturbed the etioplast-to-chloroplast transition or blocked photosynthetic electron transport, reduced light-induced plasmodesmata formation between mesophyll and bundle sheath cells of C_4_ *G. gynandra*. We conclude that chloroplast function, and in particular photosynthetic electron transport, play important roles in controlling the formation of secondary plasmodesmata in the C_4_ leaf.

The inhibitory effect of DCMU on plasmodesmata formation could be associated with signaling from a dysfunctional photosynthetic electron transport chain, or because less photosynthate is produced. To test the latter hypothesis plants were grown on sucrose during DCMU treatment. No distinguishable effects on phenotype of the seedlings or etioplast-to-chloroplast development were detected (Fig. **6a,b**) and provision of sucrose did not rescue the reduction in *F_v_/F_m_* caused by DCMU (Fig. **6c,d**). However, when we quantified plasmodesmal frequencies in a total of 1655 cell interfaces DCMU-treated seedlings supplemented with sucrose had plasmodesmal frequencies at the mesophyll-bundle sheath interface comparable to those in untreated seedlings (Fig. **6e**, p > 0.05) indicating full rescue by sucrose of the DCMU-induced inhibition of plasmodesmata formation. Thus, when photosynthetic electron transport is inhibited, sucrose is sufficient to restore plasmodesmata formation at the mesophyll-bundle sheath cell interface of *G. gynandra*.

**Figure 6.**
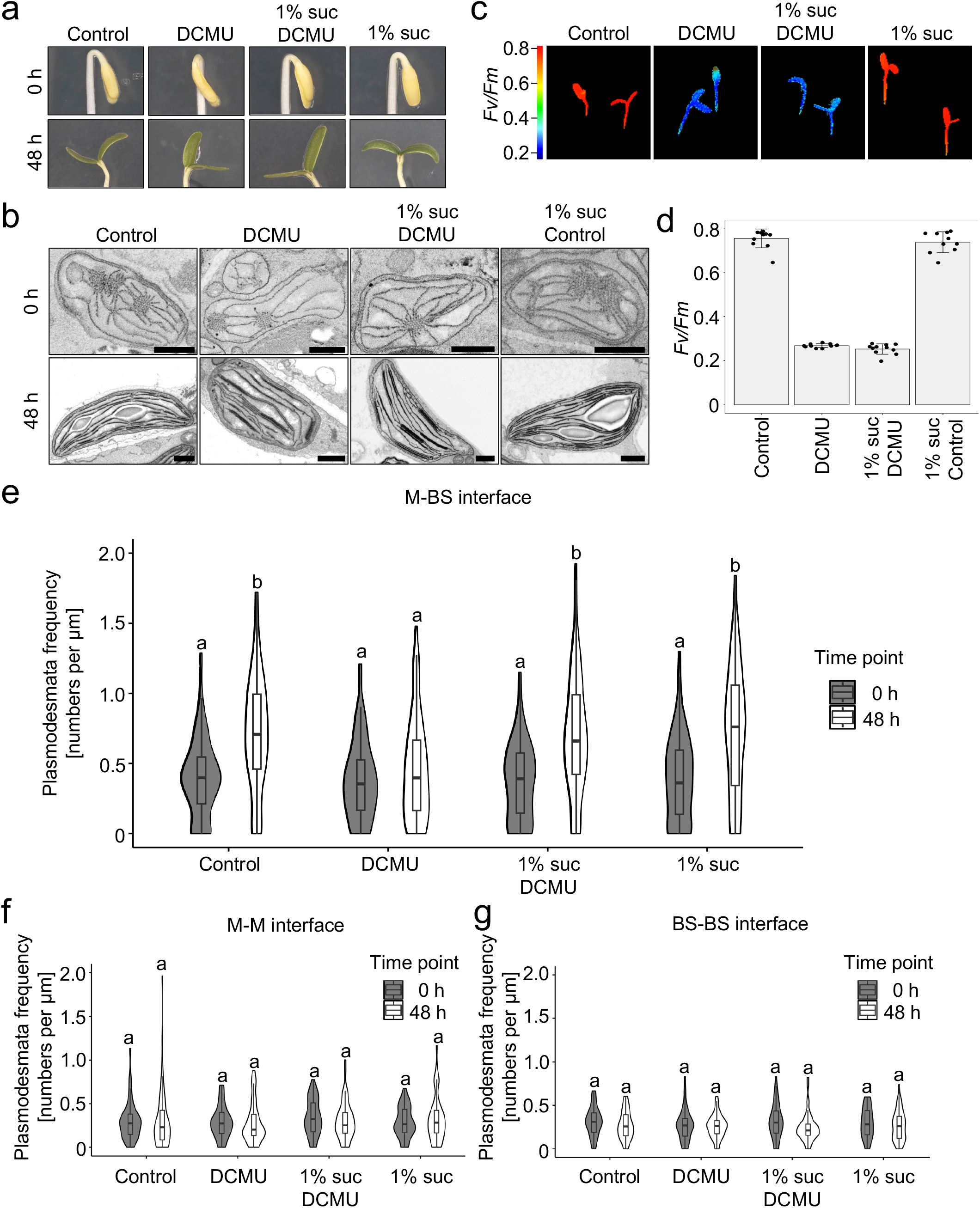
Sucrose rescues DCMU mediated inhibition of plasmodesmata formation at the M-BS interface. **(a)** Representative images of DCMU-treated *G. gynandra* seedlings during deetiolation (at 0 h and 48 h) with or without exogenous 1% (w/v) sucrose. (**b**) Scanning electron micrographs of etioplasts (0 h) and mature chloroplasts (48 h) of DCMU-treated and untreated (Control) *G. gynandra* seedlings with or without exogenous 1% (w/v) sucrose. Scale bar represents 1 μm **(c)** Chlorophyll fluorescence images of maximum quantum efficiency of PSII photochemistry (*Fv/Fm*) from 48 h deetiolated, untreated and DCMU-treated seedlings. **(d)** *Fv/Fm* measured in *G. gynandra* 48 h after light induction. Bars represent mean ± standard deviation from *n* = 7-8 individual seedlings, dots represent individual data points. (**e-g**) Plasmodesmata frequency per μm cell interfaces in *G. gynandra* cotyledons quantified during the dark to light transition (0 h and 48 h time point) and DCMU treatment, with and without additional 1% (w/v) sucrose supply using high-resolution 2D SEM maps: (**e**) M-M, (**f**) M-BS and (**g**) BS-BS. All interfaces were quantified from cotyledon samples of at least 3 individual seedlings (biological replicates) per time point. (**e**) For M-BS interface: 0h control_nosuc *n* = 96, 0h DCMU_nosuc *n* =82, 0h control_suc *n* = 84, 0h DCMU_suc *n* = 87, 48h control_nosuc *n* = 79, 48h DCMU_nosuc *n* = 98, 48h control_suc *n* = 101, 48h DCMU_suc *n* = 96 cell interfaces were quantified. (**f**) For M-M interface: 0h control_nosuc *n* = 64, 0h DCMU_nosuc *n* =57, 0h control_suc *n* = 65, 0h DCMU_suc *n* = 63, 48h control_nosuc *n* = 55, 48h DCMU_nosuc *n* = 60, 48h control_suc *n* = 58, 48h DCMU_suc *n* = 55 cell interfaces were quantified. (**g**) For BS-BS interface: 0h control_nosuc *n* = 65, 0h DCMU_nosuc *n* =62, 0h control_suc *n* = 53, 0h DCMU_suc *n* = 57, 48h control_nosuc *n* = 48, 48h DCMU_nosuc *n* = 53, 48h control_suc *n* = 62, 48h DCMU_suc *n* = 55 cell interfaces were quantified. The box and whiskers represent the 25 to 75 percentile and minimum-maximum distributions of the data. Letters show the statistical ranking, pairwise comparison of 0h and 48 h time point for each treatment, using a *post hoc* Tukey test (different letters indicate statistically significant differences at P < 0.05). Values indicated by the same letter are not statistically different.

## DISCUSSION

### Increased plasmodesmata frequency is a conserved C_4_ trait

A critical feature of the C_4_ pathway is the spatial separation of biochemical processes such that CO_2_ can be concentrated around RuBisCO. The consequence of this partitioning of photosynthesis is an absolute requirement for the exchange of metabolites between cell types. In C_4_ grasses this has long been associated with increased plasmodesmal frequency between mesophyll and bundle sheath cells (Evert et al., 1977). Previous work quantified plasmodesmata frequency at the mesophyll-bundle sheath cell interface of *G. gynandra* and yielded comparable values for plasmodesmata frequencies as in our work (Koteyeva et al., 2014), but they did not quantify plasmodesmata in any other cell interface or compared plasmodesmal frequency with related C_3_ species. Therefore, despite the very different leaf morphology between monocotyledons and dicotyledons, our results reveal that increased plasmodesmal connectivity between mesophyll and bundle sheath cells is likely a conserved trait among C_4_ plants that separate photosynthesis between two cell types. In *G. gynandra*, the mesophyll-bundle sheath interfaces had 8-13-fold higher plasmodesmata frequency than those of the closely related C_3_ species *T. hassleriana* and *A. thaliana* (Fig. **1**–**3**). This increase is comparable to plasmodesmata numbers and distributions reported for C_4_ grasses (Botha et al., 1992; Danila et al., 2016). Danila et al. (2018) further reported that C_4_ grasses running the NAD-ME subtype of C_4_ photosynthesis had the highest numbers of plasmodesmata between mesophyll and bundle sheath cells. As *G. gynandra* also primarily uses NAD-ME to decarboxylate CO_2_ in the bundle sheath, broader analysis of C_4_ dicotyledons is required to determine the extent to which plasmodesmal frequencies correlate with the various biochemical sub-types.

Plasmodesmal frequencies at the mesophyll-bundle sheath interface of *G. gynandra* are consistent with those reported previously in this species where no analysis of closely related C_3_ plants were possible (Koteyeva et al., 2014). By quantifying plasmodesmata at all interface types and comparing plasmodesmal frequency with phylogenetically proximate C_3_ plants we demonstrate that plasmodesmata numbers are generally higher at all three types of cell interface (mesophyll-bundle sheath, mesophyll-mesophyll, bundle sheath-bundle sheath) in C_4_ *G. gynandra*. This is consistent with previous work that observed increased plasmodesmata frequencies between photosynthetic leaf cells in C_4_ grasses compared with C_3_ grasses (Danila et al., 2016).

Compared with C_4_ grasses, a distinguishing feature of the increased plasmodesmal frequency between mesophyll and bundle sheath cells of *G. gynandra* is that the increase was not associated with any detectable increase in pitfield area compared with C_3_ *T. hassleriana* or C_3_ *A. thaliana* (Supporting Information Fig. **2**). This suggests that the primary mechanism for increased plasmodesmata numbers in *G. gynandra* is an increase in pitfields. Since we were not able to visualise individual plasmodesmata within pitfields, we cannot rule out that there is a higher frequency of individual plasmodesmata number within pitfields. However, we consider this is unlikely since pitfield appearance was largely similar between *G. gynandra, T. hassleriana* and *A. thaliana* (Supporting Information Fig. **2**). In contrast, increased plasmodesmal frequency in C_4_ grasses was accompanied by increases in pitfield area such that were up to 5 times greater than those in C_3_ species (Danila et al., 2016, 2018). This difference suggests that the mechanisms by which increased plasmodesmata numbers between mesophyll and bundle sheath cells can vary between C_4_ lineages.

Flux of metabolites between cells is likely to be determined by plasmodesmata number as increased numbers can facilitate greater flux. However, bundle sheath cells are not air-tight and plasmodesmata could also contribute to CO_2_ leakiness such that a proportion of the CO_2_ concentrated in the bundle sheath diffuses back to the mesophyll. CO_2_ leakiness particularly increases during photosynthetic induction in NADP-ME type C_4_ plants such as sorghum and maize (Wang et al., 2022). Thus, it is possible that plasmodesmata number and distribution need to be optimised to allow maximum photosynthetic efficiency in C_4_ plants. Being able to accurately quantify plasmodesmal traits in diverse C_4_ species may be crucial to develop further understanding in this area, and in particular in modelling metabolite flux through the C_4_ pathway (Danila et al., 2016; Von Caemmerer, 2021). This could incorporate recent models of metabolite diffusion through plasmodesmata such as the geometric and narrow escape models (Denim et al., 2019; Hughes et al., 2021).

### Light triggers rapid plasmodesmata formation in pre-existing cell walls

In C_4_ grasses the developmental cue that enhances plasmodesmata formation between mesophyll and bundle sheath cells is not known. However, *Setaria viridis* and maize show some plasticity in plasmodesmal density in response to growth irradiance (Danila et al., 2019). Our data provides a direct link between light and photosynthesis in establishing plasmodesmal frequency by showing that light rapidly triggers the formation of plasmodesmata at the mesophyll-bundle sheath interface in *G. gynandra*.

Plasmodesmata are either formed *de novo* during cell division by trapping ER strands between enlarging Golgi-derived vesicles in new cell walls (primary plasmodesmata) or formed in pre-existing cell walls (secondary plasmodesmata) (Hepler, 1982; Ehlers and Kollmann, 2001; Faulkner et al., 2008). We believe that the increase in plasmodesmata numbers between mesophyll and bundle sheath cell during dark to light transition is primarily driven by the formation of secondary plasmodesmata for the following reasons. Firstly, cotyledon growth from dark to light is thought to be exclusively driven by cell expansion and not cell division in Arabidopsis (Tsukaya et al., 1994; Stoynova-Bakalova et al., 2004). Secondly, the basic structure of bundle sheath cells was already formed in dark grown seedlings, and the formation of plasmodesmata was rapid. Our SEM mapping technique provided sufficient resolution to observe branching in plasmodesmata (Fig. **2**,**4–6**), but interestingly we did not observe any structural differences between the plasmodesmata in different cell interfaces. Although primary and secondary plasmodesmata can be sometimes distinguished by structure, where secondary plasmodesmata are more branched, this is highly dependent on other factors such as leaf age and sink-source transition (Roberts et al., 2001).

### A role for metabolism and organelles in formation of plasmodesmata

Our results suggest that chloroplasts, and more specifically photosynthesis, fuel the formation of secondary plasmodesmata between mesophyll and bundle sheath cells in C_4_ *G. gynandra*. Inhibition of photosynthesis and chloroplast development through the application of inhibitors greatly reduced plasmodesmata formation during deetiolation but this effect could be rescued by the exogenous supply of sucrose (Fig. **5****,6**). To our knowledge, a role of photosynthate in controlling formation of plasmodesmata has not been proposed previously. However, some findings are consistent with this hypothesis. For example, in rice constitutive overexpression of the C_4_ maize *GOLDEN2-LIKE* transcription that controls chloroplast biogenesis (Waters et al., 2008) not only activated chloroplast and mitochondria development in bundle sheath cells but also increased plasmodesmata numbers (Wang et al., 2017). Moreover, in *A. thaliana* links between organelles and plasmodesmata have been reported. *A. thaliana* mutants with altered cell-to-cell connectivity and/or plasmodesmata structure such INCREASED SIZE EXCLUSION LIMIT1 and 2 (ISE1/ISE2) encode mitochondrial and chloroplast RNA helicases respectively (Kobayashi et al., 2007; Stonebloom et al., 2009), while the GFP ARRESTED TRAFFICKING1 (GAT1) locus encodes a chloroplast thioredoxin (Benitez-Alfonso et al., 2009). However, the mechanisms of how these organelle-localized proteins affect plasmodesmata formation are not understood. Retrograde signaling from chloroplast to nucleus has also been proposed to control plasmodesmata formation and regulation (Burch-Smith et al., 2011; Ganusova et al., 2020). The fact that exogenous supply of sucrose is sufficient to sustain plasmodesmata formation in the presence of DCMU (Fig. **6**) strongly suggests a direct metabolic role of chloroplasts in the enhanced formation of plasmodesmata in the C_4_ leaf. This may involve sucrose or photosynthesis providing energy required for plasmodesmata formation, or sucrose acting as a signalling molecule to trigger plasmodesmata formation via sugar signalling. Further work is required to address how sugar controls plasmodesmata formation in *G. gynandra*. In summary, our work demonstrates that increased plasmodesmal connectivity is likely conserved trait found in both C_4_ dicotyledons and monocotyledons. Moreover, the enhanced formation of plasmodesmata between mesophyll and bundle sheath cells of C_4_ leaves is coordinated and dependent on photosynthesis. Evolution therefore appears to have wired the enhanced formation of plasmodesmata in C_4_ leaves to the development of chloroplasts and432 ultimately the induction of photosynthesis.

## DATA AVAILABILITY

The data supporting the findings of this study are available from the corresponding author upon request.

## ACKNOWLEDGEMENTS

The work was funded by the Advanced European Research Council (grant 694733 REVOLUTION to J.M.H.). T.B.S. was supported by the Swiss National Science Foundation (SNSF) Early Postdoc Mobility Fellowship (P2EZP3_181620), the SNSF Postdoc Mobility Fellowship (P500PB_203128) and the EMBO Long-Term Fellowship (ALTF 531-2019). C.F. was funded by the European Research Council (grant 725459 INTERCELLAR to C.F.) and Biotechnology and Biological Research Council Institute Strategic Programme (Plant Health, BBS/E/J/000PR9796 to the John Innes Centre). For the purpose of open access, the authors have applied a Creative Commons Attribution (CC BY) license to any Author Accepted Manuscript version arising from this submission. We thank Filomena Gallo and Lyn Carter from the Cambridge Advanced Imaging Centre for the electron microscopy sample preparation as well as the support during the image acquisition. We thank Miriam Lucas from Scope M at ETH Zurich for advice and assistance with SBF-SEM. We also thank for Zhengao Di for help with the chlorophyll fluorescence measurement settings.

## AUTHOR CONTRIBUTIONS

T.B.S. and J.M.H. conceived and directed the research; T.B.S, C.F., S.C.Z. and J.M.H. designed the experiments; T.B.S., K.M. and S.E. performed the research and T.B.S. analyzed the data; T.B.S. and J.M.H. wrote the article with input from all the authors.

## SUPPORTING INFORMATION

**Supporting Information Video 1. Compiled video of sequential 50 nm sections of M-BS cell interface in mature leaves of C_4_ *G. gynandra*.**

**Supporting Information Video 2. Compiled video of sequential 50 nm sections of M-BS cell interface in mature leaves of C_3_ *T. hassleriana*.**

**Supporting Information Video 3. Compiled video of sequential 50 nm sections of M-BS cell interface in mature leaves of C_3_ *A. thaliana*.**

**Supporting Information Figure 1. Transmission electron micrographs of M-BS, M-M and BS-BS cell interfaces.**

**Supporting Information Figure 2. Comparison of plasmodesmal frequencies quantified using 2D SEM and 3D SBF-SEM.**

**Supporting Information Figure 3. Pitfield area is not increased in C_4_ *G. gynandra* compared to C_3_ *A. thaliana* and *T. hassleriana*.**

**Supporting Information Figure 4. Extended dark treatment for 48 h does not increase plasmodesmata frequency in *G. gynandra* cotyledons.**

**Supporting Information 5. Chloroplast inhibitors have limited effect on light-induced cotyledon expansion, but affect plasmodesmata formation.**

