## Supporting Information Figures for "Plasmodesmal connectivity in C_4_ *Gynandropsis gynandra* is induced by light and dependent on photosynthesis"

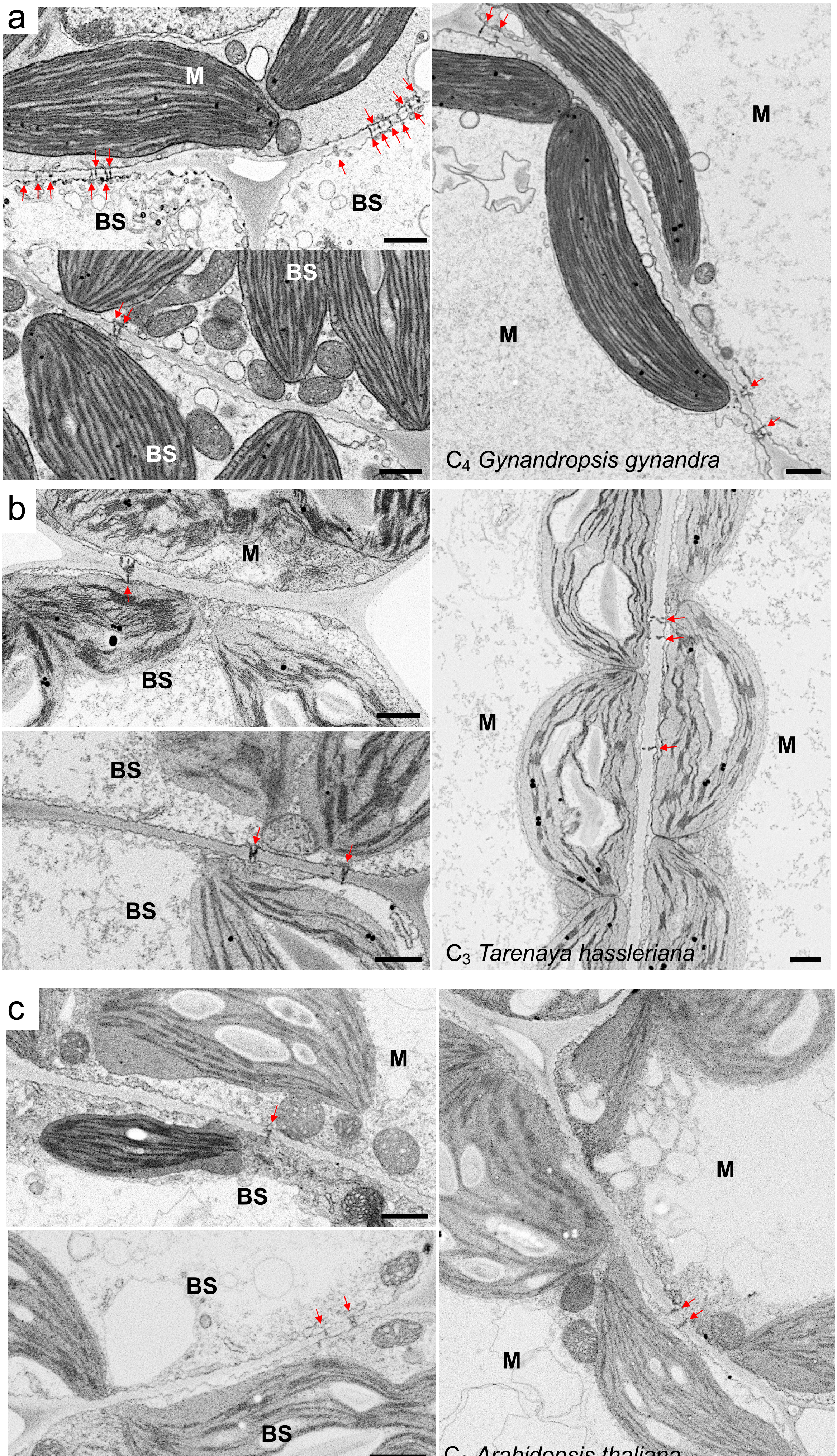

a
M
BS
BS
BS
BS
M
M
C4 Gynandropsis gynandra
b
M
BS
BS
BS
M
M
C3 Tarenaya hassleriana
c
M
BS
BS
BS
M
M
C3 Arabidopsis thaliana

**Supporting Information Figure 1. Transmission electron micrographs of M-BS, M-M and BS-BS cell interfaces.** Representative interfaces in (**a**) $C_4$ *G. gynandra*, (**b**) $C_3$ *T. hassleriana* and (**c**) $C_3$ *A. thaliana* leaves. Mature leaves were harvested from 4-week-old *G. gynandra* and *T. hassleriana* plants and 3-week-old *A. thaliana*. Red arrows indicate individual plasmodesma. Scale bar = 1 µm

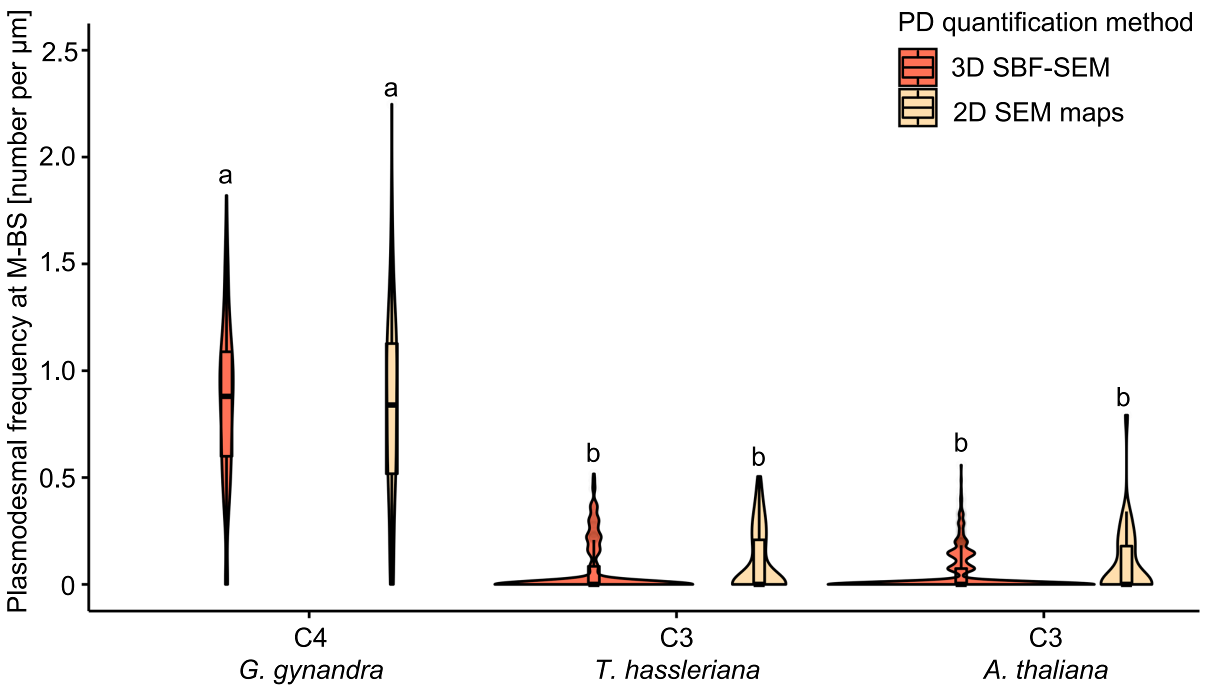


**Supporting Information Figure 2. Comparison of plasmodesmal frequencies quantified using 2D SEM and 3D SBF-SEM.** Numerous M-BS cell interfaces were quantified in 2D SEM maps, while few M-BS cell interfaces were analysed in-depth using 3D SBF-SEM. The box and whiskers represent the 25 to 75 percentile and minimum-maximum distributions of the data. Letters show the statistical ranking using a *post hoc* Tukey test (different letters indicate significant differences at $P < 0.05$). Values indicated by the same letter are not statistically different.

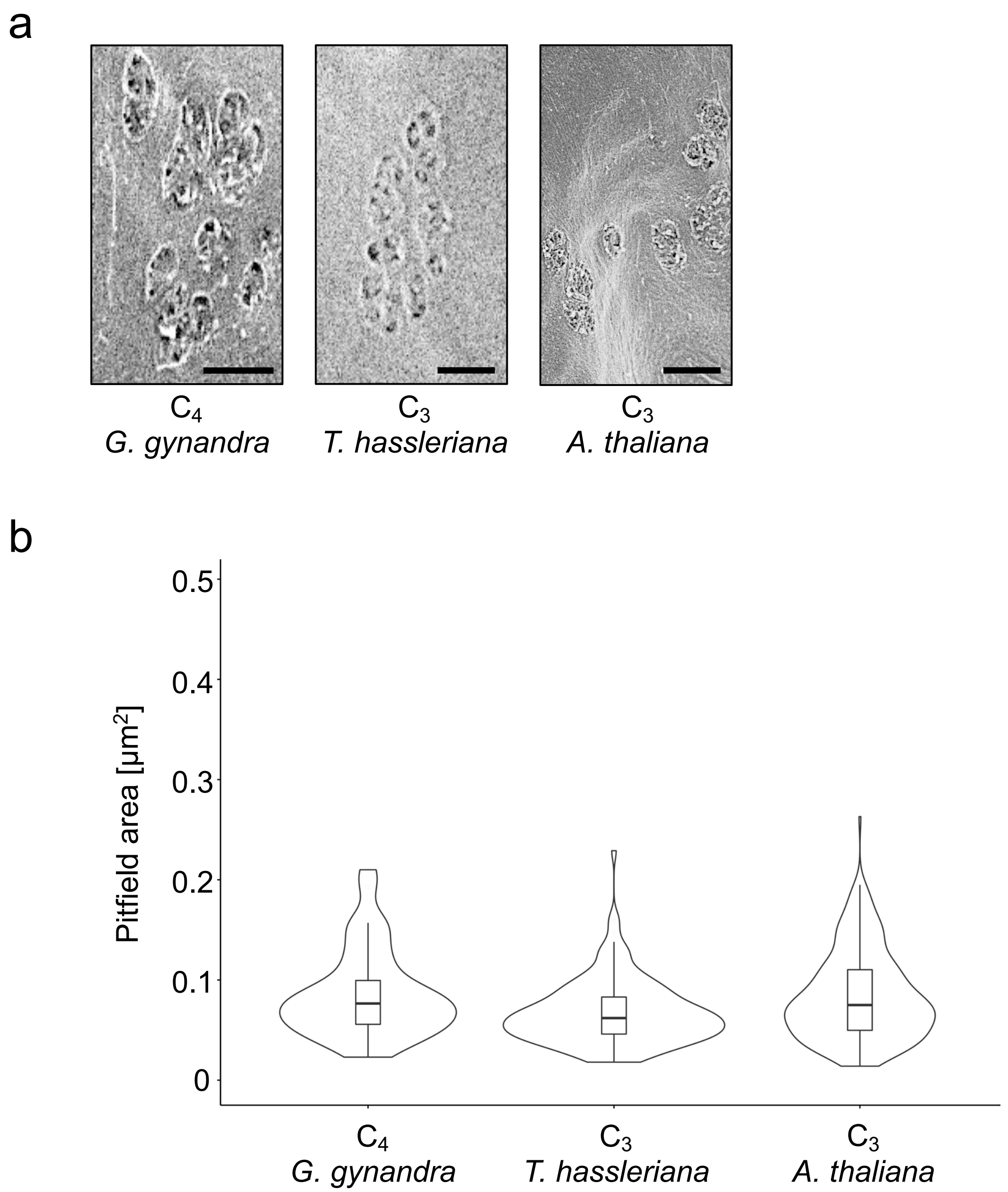


**Supporting Information Figure 3. Pitfield area is not increased in $C_4$ *G. gynandra* compared to $C_3$ *A. thaliana* and *T. hassleriana*.** (**a**) Representative scanning electron micrographs of plasmodesmata pitfields at M-BS cell interface in each species. Scale bars = 0.5 µm (**b**) Measured pitfield areas of *G. gynandra* (*n* = 260), *T. hassleriana* (*n* = 151) and *A. thaliana* (*n* = 40).

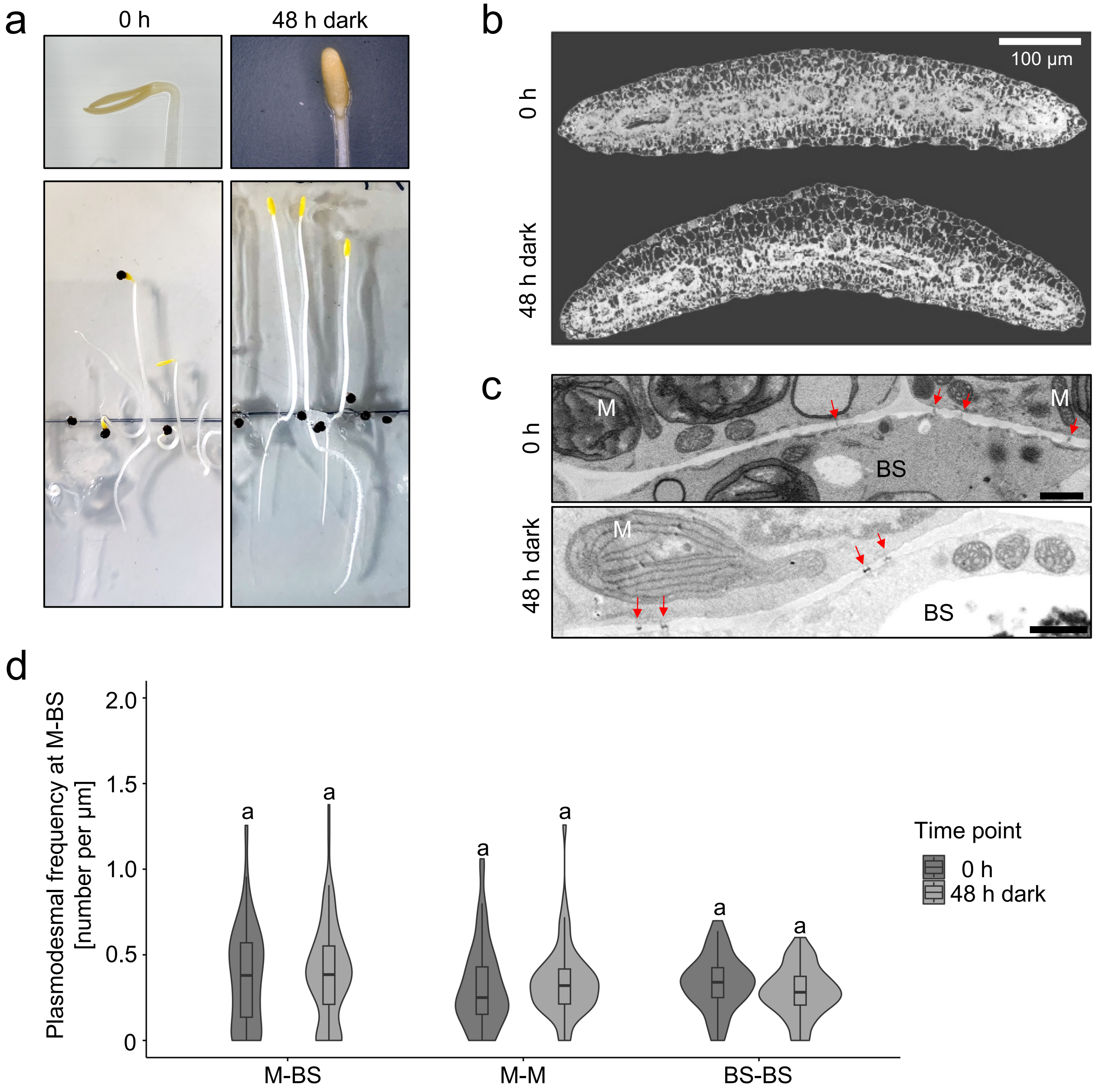


**Supporting Information 4. Extended dark treatment for 48 h does not increase plasmodesmata frequency in *G. gynandra* cotyledons.** (**a**) Photographs of 3-day-old (0) and 5-day-old (48 h extended dark) dark-grown *G. gynandra* seedlings on half-strength MS media. (**b**) Scanning electron micrographs of entire cotyledon cross sections of 0 h and 48 h extended dark-treated *G. gynandra* seedlings. (**c**) Representative scanning electron micrographs of M-BS interfaces of 0 h and 48 h extended dark-treated *G. gynandra* cotyledons. Red arrows indicate individual plasmodesma. Scale bar = 1 µm (**d**) Plasmodesmal frequency per µm cell interfaces (for M-BS, M-M and BS-BS) in *G. gynandra* cotyledons was quantified before and after extended dark treatment (0 h and 48 h dark time point) using high-resolution 2D SEM maps. For the 0 h time point, *n* = 84 (M-BS), *n* = 60 (M-M), and *n* = 41 (BS-BS) cell interfaces were quantified. For the 48h dark time point, *n* = 91 (M-BS), *n* = 60 (M-M), and *n* = 58 (BS-BS) cell interfaces were quantified. All interfaces were quantified from cotyledon samples of at least 3 individual seedlings (biological replicates) per time point. The box and whiskers represent the 25 to 75 percentile and minimum-maximum distributions of the data. Letters show the statistical ranking using a *post hoc* Tukey test (different letters indicate significant differences at P<0.05). Values indicated by the same letter are not statistically different.

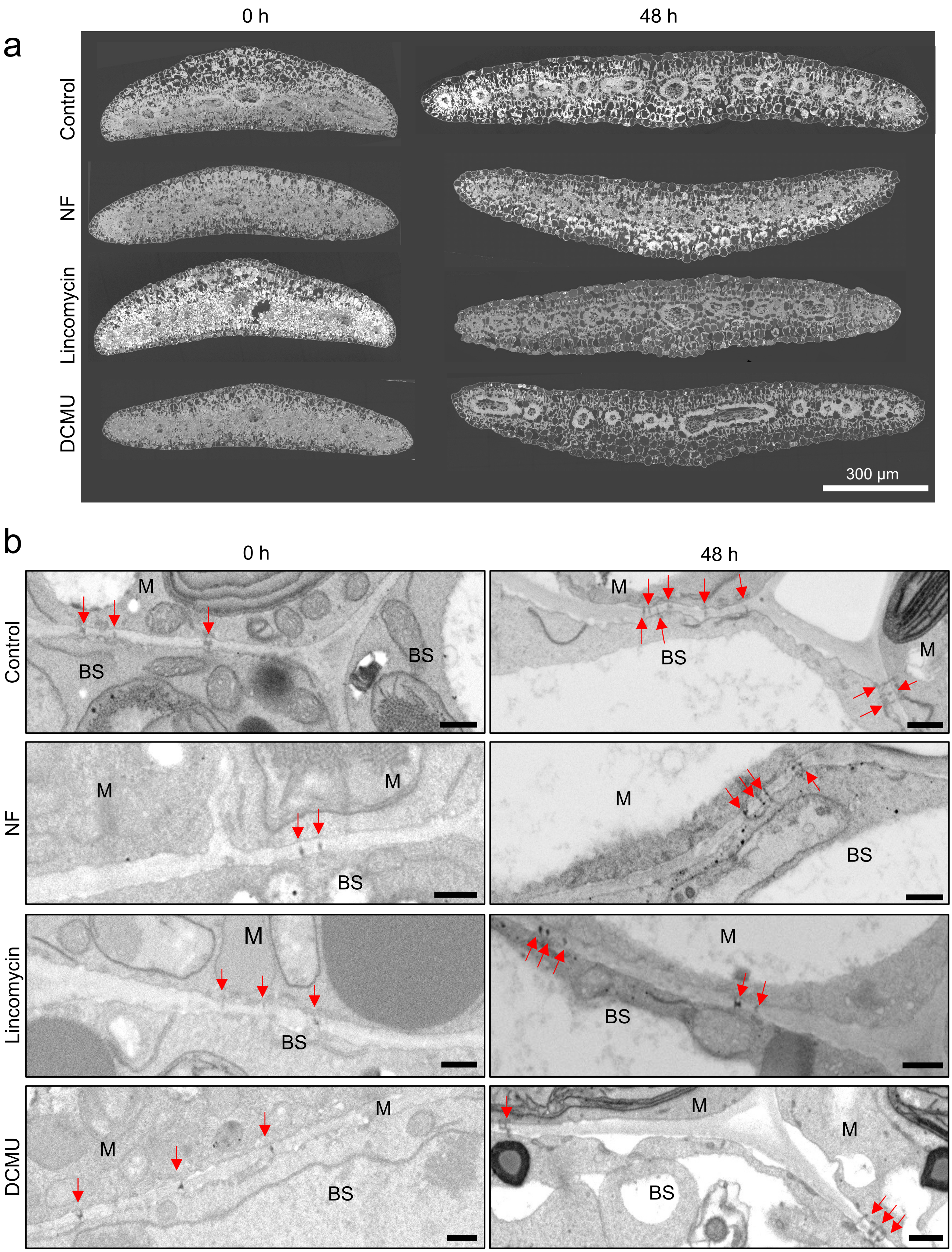
a
0 h
48 h
Control
NF
Lincomycin
DCMU
300 μm
b
0 h
48 h
Control
NF
Lincomycin
DCMU
M
BS

**Supporting Information 5. Chloroplast inhibitors have limited effect on light-induced cotyledon expansion, but affect plasmodesmata formation.** The effect of norflurazon (NF), lincomycin (Linco) and DCMU were tested. (**a**) Scanning electron micrographs of entire cotyledon cross sections of *G. gynandra* seedlings, 0 h and 48 h after light induction. (**b**) Representative scanning electron micrographs of M-BS interfaces of 0 h and 48 h during de-etiolation of *G. gynandra* cotyledons treated with NF, Linco and DCMU, as well as untreated seedlings (Control). Red arrows indicate individual plasmodesma. Scale bar = 1 µm
